# Ultrasound-Induced Reorientation for Multi-Angle Optical Coherence Tomography

**DOI:** 10.1101/2023.10.04.560819

**Authors:** Mia Kvåle Løvmo, Shiyu Deng, Simon Moser, Rainer Leitgeb, Wolfgang Drexler, Monika Ritsch-Marte

## Abstract

Organoid and spheroid technology have recently provided great insights into oncology, developmental biology as well as personalized medicine. Among the methods to optically monitor the structural and functional organization of such samples, optical coherence tomography (OCT) has emerged as an excellent, label-free approach. Mature organoids, however, are often too opaque for OCT due to regions of strong attenuation. This leads to severe artifacts and reduced morphological tissue information in the reconstruction, since the far-side of the specimen is not reachable. Access to multi-angle views of OCT is therefore highly desirable. This aligns with another problem affecting certain goals of organoid research: The sample needs to be embedded in a growth scaffold such as Matrigel, whereas freely floating objects would not suffer from confinement and be more easily accessible for mechanical or chemical probing. Here we present ULTrasound-Induced reorientation for Multi-Angle-OCT (ULTIMA-OCT), a solution overcoming these limitations. By inserting a small 3D-printed acoustic trap to a spectral-domain OCT system, acoustic actuation enables contact-free levitation and finely tunable stepwise reorientation of samples such as zebrafish larvae and tumor spheroids, in a controlled and reproducible manner. This enables tomographic reconstruction of (sub-)mm samples with enhanced penetration depth and reduced attenuation artifacts, by means of a model-based algorithm we developed. We show that this approach is able to fuse the diverse multi-angle OCT volumes for a joint recovery of 3D-reconstruction of reflectivity, attenuation, refractive index and position registration for zebrafish larvae. We believe that our approach represents a powerful enabling tool for developmental biology and organoid research.

## INTRODUCTION/MOTIVATION

A steep increase in organoid and spheroid research could be witnessed in recent years, providing vital insights into developmental biology and oncology. A strong motivation for this is the potential of organoids and cancer spheroids to reduce animal experimentation to some extent [1]. Spheroids can be grown with the support of an extracellular matrix or scaffold-free [2]. Organoids are usually grown in Matrigel, which is derived from the secretion of a type of mouse sarcoma cells. Matrigel is complex and variable which gives rise to a certain irreproducibility. Moreover, it is known that the matrix scaffolds have a mechanical impact which is often not well understood or characterized [3]. Therefore, in the past few years considerable effort has been directed towards Matrigel-free organoid growth [4].

In response to such problems, contact-free levitation of samples has been sought. On the single-cell level, it is possible to use holographic optical tweezers, but for cell-clusters reaching mm-size, optical trapping would lead to over-heating due to the intensities needed to counteract the growing weight [5], even in the most favorite case of counter-propagating optical beams creating a trapping region *between* two laser spots, as in the macro-tweezers system [6], [7]. To tackle larger biological samples, ultrasound techniques for levitation and actuated handling have been developed: Standing bulk acoustic waves (BAW) operating at (sub-)MHz frequencies push biological cells into low–pressure regions (planes, lines or spots depending on the number of orthogonal standing waves) [8–10], and by modulation of the acoustic waves objects can be transiently or continuously rotated [11– 15]. Surface acoustic waves (SAW) can also be used, e.g., to create acoustofluidic rotational tweezers for morphological phenotyping of zebrafish larvae [16]. A big advantage of acoustic levitation is the scaffold-free confinement, which is more open to perform various assays, such as irradiating parts by light, adding chemicals and pharmaceuticals by micro-fluidics, mechanical probing with tips, and other assays.

Optical Coherence Tomography (OCT) [17–19], a technique based on low coherence interferometry, can reconstruct micrometer sample morphology from the backscattered light with high imaging speed (MHz A-scan rate) and has been widely employed for (bio)medical applications. Optical Coherence Microscopy (OCM), which uses higher numerical aperture (NA), can achieve high lateral resolution but usually is limited in penetration depth. OCT/OCM recently has emerged as a valuable imaging modality for organoids and spheroids [20–22], providing high-resolution information on the internal structural organization inside the organoids non-invasively and lable-free.

Nevertheless, mature specimens often become optically dense, intractable not only for Optical Diffraction Tomography (ODT) [23, 24] but also for OCT, leading to shadows and limited tissue morphology information. Shadow removal algorithms have been developed for OCT images of the optic nerve head [25–27]. However, removing the shadows cast by high-attenuation structures like the eye of a wild-type zebrafish larva remains challenging, because the OCT incident light can be fully occluded. 3D optical coherence refraction tomography [28] compensated for this issue by con-trolling the incident beam angle and position using a parabolic mirror, but it was limited to*±* 75° angular orientation and needed to immobilize samples like zebrafish embryos in agarose gel.

In this work, we present an easy-to-use solution overcoming the above explained limitations and problems encountered when imaging organoids, spheroids or developing organisms: ULTrasound-Induced reorientation for Multi-Angle OCT (ULTIMA OCT) uses a small add-on microfluidic chamber with tunable acoustic waves to stably and reproducibly rotate the cell cluster into several orientations. This enables OCT imaging from different viewing angles in the full range of 360° around the sample’s major axis, which makes 3D tomographic reconstruction feasible also for optically dense samples. The immobilization of the sample is contact-free and does not involve any rotating mechanical parts, nor any elements obstructing optical imaging or introducing optical aberrations.

The price to pay in this approach is the fact that one cannot precisely choose the exact viewing angles, since the stable trapping positions and orientations in the acoustic force fields to some extent also depend on the unknown sample itself. However, we provide a generally applicable solution, i.e. a model-based algorithm which can deal with the added complexity of tomographic reconstruction with viewing angles that are not precisely known *a priori*.

## RESULTS

### Working principle of ULTIMA-OCT

The workflow of ULTIMA-OCT and its ingredients are explained schematically in Fig. 1, and a schematic animation of the data acquisition procedure can be found in Supplementary Movie 1. Acoustic radiation forces are used to levitate the sample and to induce transient rotations in a fluid chamber. Each new orientation is a stable trapping position, and we perform OCM scanning of the sample at a desired number of orientation angles. The 3D OCM data is then processed by a compressive model-based algorithm to extract the underlying reflectivity map, while also yielding information about the attenuation and refractive index (RI) contrast maps as well as the position and orientation parameters of the trapped sample.

**Figure 1.**
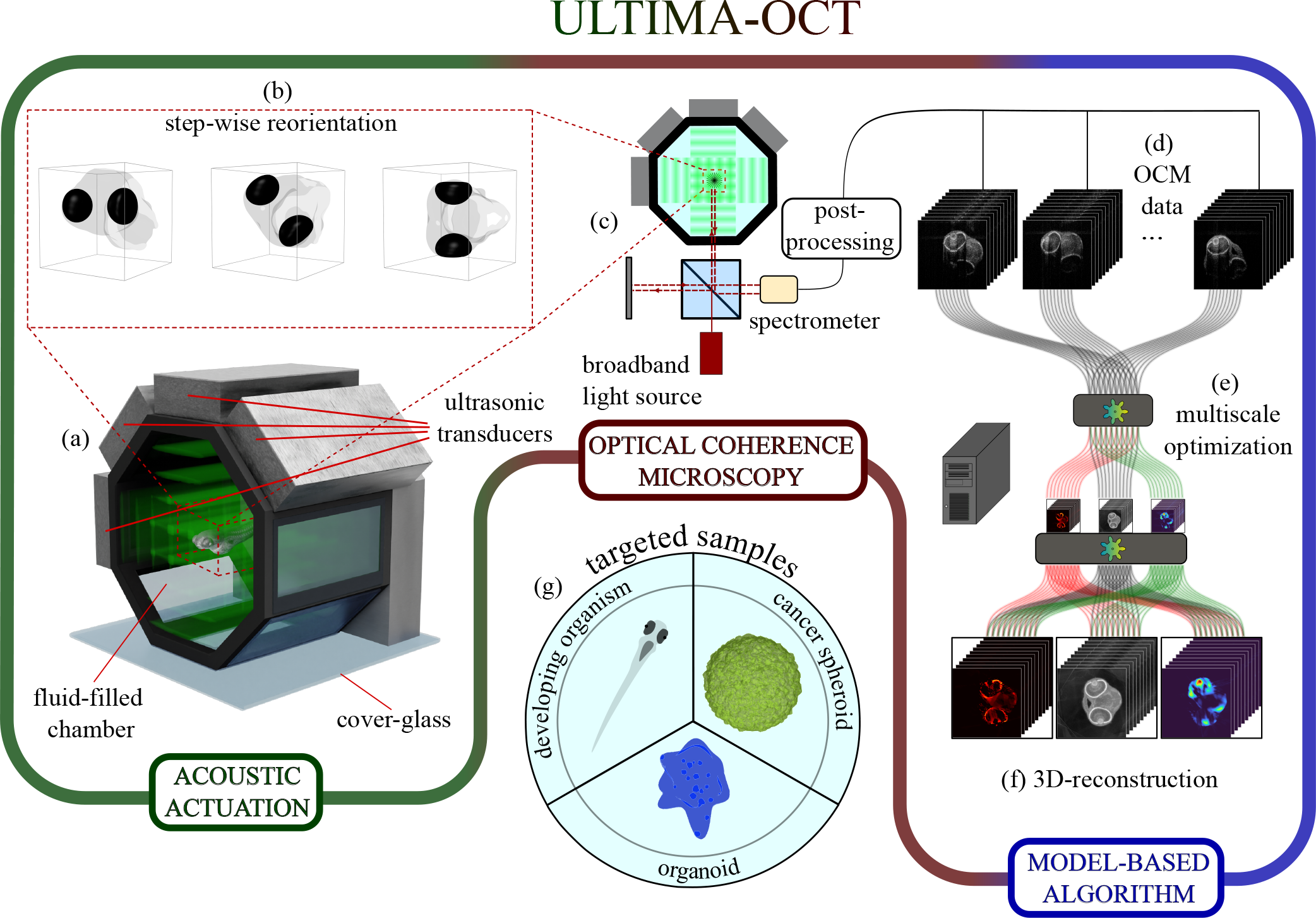
Schematics and workflow of ULTrasound-Induced reorientation for Multi-Angle-OCT (ULTIMA-OCT). (a) depicts the fluid-filled acoustic chamber, in which the sample is levitated and reoriented by means of acoustic actuation. The specimen is step-wise rotated into several stable trapping positions (b), and optical coherence microscopy (OCM) imaging is performed in each of them (c). The acquired OCM data is postprocessed (d) and fed into a multiscale optimization algorithm (e), which performs fusion of the images and outputs 3D reconstructions of reflectivity, attenuation and refractive index (RI) maps. In (g) an exemplary collection of samples is shown, where ULTIMA-OCT can be applied.

### Acoustic actuation

The acoustic manipulation chamber consists of a 3D printed frame with a symmetric octagon cross-section with 4 piezo-electric plate transducers and 4 reflectors around its sides (Fig. 2 and Supplementary Fig. 1). We apply an AC signal to the transducers to propagate bulk acoustic waves (BAWs) in 4 directions in the liquid-filled chamber and generate acoustic standing waves in each direction upon reflection. The standing waves have pressure nodes every *λ/*2 (*≈*1 mm around 700 kHz) along each propagation direction in the fluid chamber. The front and back of the octagon frame is sealed with a cover-slip and an aluminium plate respectively, and imaging is performed through the bottom cover-slip that also acts as the reflector for the acoustic waves from the top-transducer. Our current device is developed to accommodate a range of sample sizes and shapes and we have demonstrated this by manipulating samples, from highly asymmetric mm-sized 3-5 days post fertilization (dpf) zebrafish embryos to less asymmetrically shaped sub-mm-sized melanoma spheroids. We rely on the resonant enhancement of the waves in the fluid chamber to get sufficient forces to levitate our targeted biological samples. The specific chamber dimensions used for the results presented here were found based on numerical simulations and experimental verification (see “Methods” and Supplementary Methods for details on chamber dimensions and acoustic resonances). Levitation by the top transducer was achieved at a minimum peak-to-peak driving voltage of 20 V, corresponding to a maximum pressure amplitude of 80 kPa (see Supplementary Methods).

**Figure 2.**
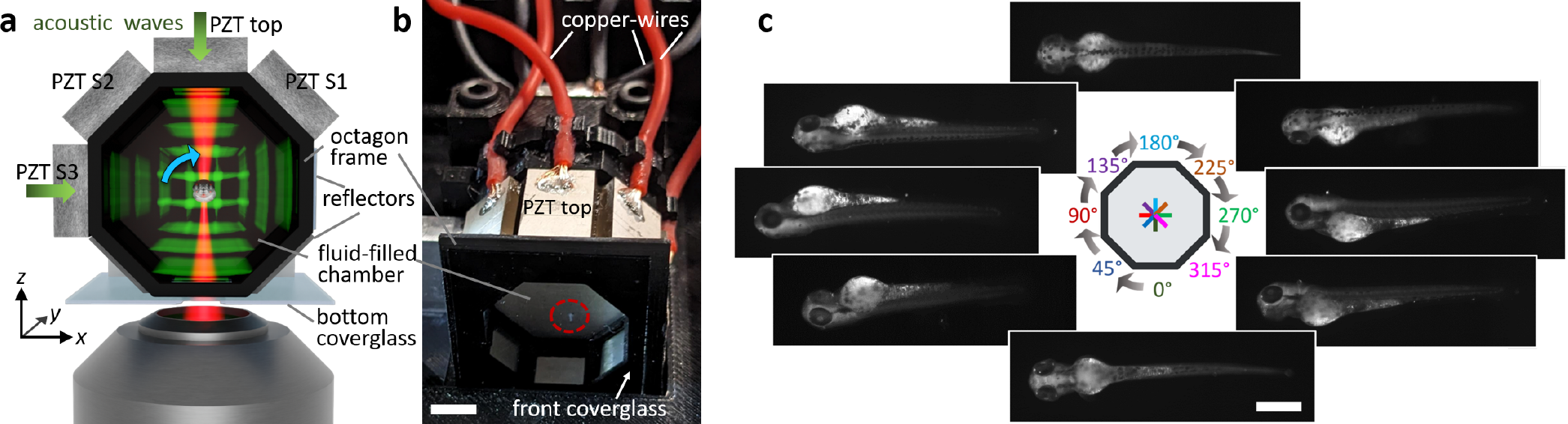
**Acoustic actuation for multi-angle imaging. a** Illustration of acoustic manipulation of levitated zebrafish embryo (not to scale). By coupling bulk acoustic waves into the fluid-filled chamber from multiple directions, acoustic standing waves (green) are generated upon reflection, to levitate the sample and induce transient rotations for optical imaging (red beam), e.g. for multi-angle high-speed OCT through the bottom cover-glass of the 3D printed octagon frame (black). The direction of rotation in the *x-z* -planes is indicated (blue arrow). **b** Assembled octagon chamber with levitated zebrafish embryo (inside stippled red circle), scale bar: 5 mm. **c** The optimization of the acoustic actuation can be carried out on an inverted microscope with optical image acquisition. As an example of darkfield (oblique illumination) images of a wild-type 3 dpf zebrafish embryo are shown here, for a selection of 8 chosen angles of acoustic reorientation, scale bars: 600 µm.

To characterize and optimize our contact-less trapping platform for reorientation and multi-angle image acquisition, we used fixated 3 dpf zebrafish embryos, as they are readily available samples that are perfectly suited to demonstrate the benefits of our approach. To observe the zebrafish while tuning the acoustic settings for stable reorientation, we used an inverted microscope with oblique illumination through the front cover-glass of the chamber, acquiring dark-field images (Fig. 2**c** and details in Supplementary Methods). With the acoustic radiation forces [29–32] from the top transducer, we levitate the sample against gravity in one of the nodal planes in the center region of the chamber where all 4 acoustic waves intersect. With the additional radiation forces from one side-transducer, we align the sample with its major axis to the length of the chamber (*y* -axis in Fig. 2).

By changing the voltage on the transducers, and hence the relative magnitudes of the acoustic radiation forces in each direction, we generate an acoustic restoring torque [9, 33, 34] acting on the sample. The torque direction is perpendicular to the acoustic propagation directions, hence parallel to the *y* -axis and the direction of rotation is in the *xz* -planes (Fig. 2). With a dominating top transducer, the sample is aligned with its minor axis to the steepest trap-stiffness in *z* -direction (90° in Fig. 2c). When we increase the amplitude of one of the side transducers in a step-wise manner, we rotate the pressure landscape and the sample is rotated in a step-wise fashion until the sample is aligned with its minor axis to the now dominating forces from the side-transducer. We alternate between increasing and decreasing the amplitudes between pairs of the transducers in a sequence to rotate the sample 360° about its major axis, while ensuring levitation (top transducer voltage is tuned, but never zero).

Moreover, we found that for sufficiently asymmetric samples one can precisely control the orientation in a more efficient way, by choosing two orthogonal modes of exactly the same frequency in the chamber (by e.g., the top transducer and the orthogonal side-transducer S3 in Fig. 2) and adjusting the relative amplitude and phase (see Supplementary Methods for details on acoustic actuation), similar to [11] but with additional levitation in our upright (not horizontal) chamber. In Fig. 2c we show dark-field images (2 stitched tiles per image) from 8 different orientations of a 3 dpf zebrafish, but we can reorient this sample with a much finer step-size, see Supplementary Movie 2 showing reorientation by two transducers of a zebrafish embryo. These results imply that, for certain types of samples, one could potentially use chambers with a square cross-section and only two orthogonal transducers for reorientation around one axis.

To demonstrate our capability of extending the outlined acoustic manipulation to other types of samples, we also trapped and reoriented melanoma spheroids (see Supplementary Fig.4). These samples were smaller and less asymmetric than the zebrafish embryos, but we could reorient it around its major axis by the same-frequency two-transducer actuation described above. At certain settings, however, we induce a sustained rotation of the sample instead of a transient rotation. We can change the rotation direction by changing the relative phase by 180° between the two orthogonal transducers inducing the acoustic spinning torque [12, 13] (see Supplementary Movie 3). It has so far always been possible to avoid these settings and stably reorient such samples 360° around one axis. In each orientation the sample is held stably without any significant motion, see Supplementary Movie 2 and 3. Our trapping platform can accommodate a large range of sample sizes and shapes, but each new sample needs its own fine-tuning of the acoustic settings, which is performed on the fly while directly observing the object.

### High-resolution OCM imaging

To verify the feasibility of ULTIMA-OCT, biological samples such as fixated zebrafish larvae and melanoma spheroids were tested. The OCM system successfully captured the 3D data of the samples levitated and reoriented in the acoustic chamber. Fig. 3 shows OCM images of a 3 dpf less-pigmented *Mitfa*^*b*692*/b*692^/*ednrb1*^*b*140*/b*140^ zebrafish and a melanoma spheroid, respectively, imaged from one direction.

**Figure 3.**
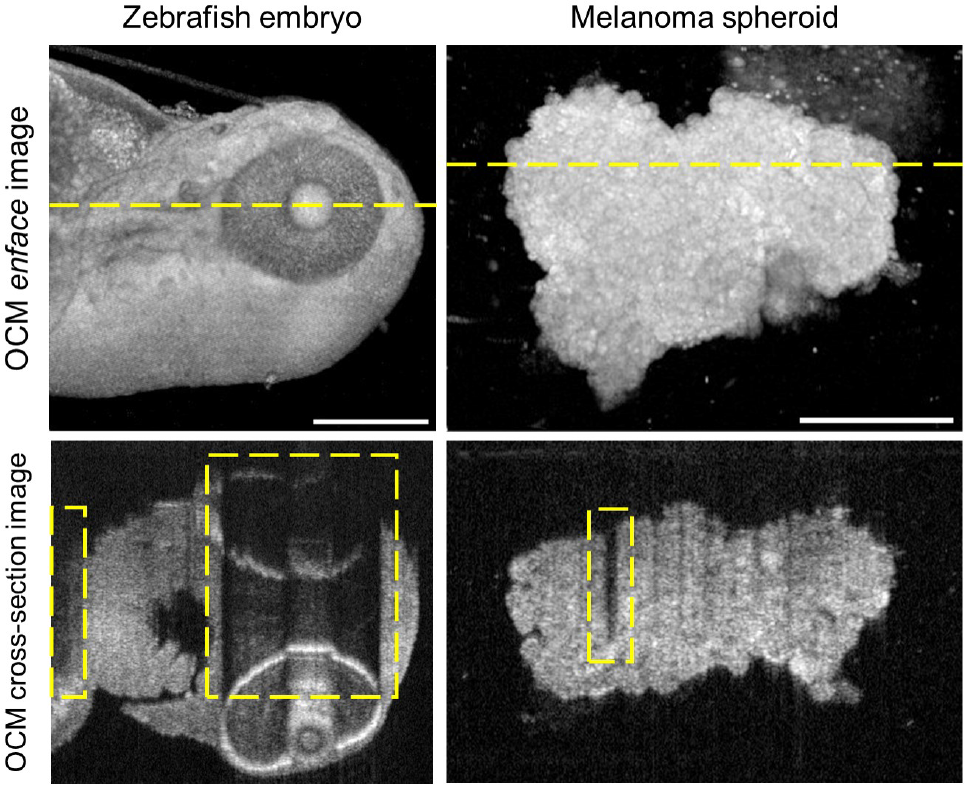
OCM limitations for imaging from only one direction. Cross-section image positions are indicated by the yellow dash lines in the OCM *en face* images (average intensity projection for zebrafish embryo and standard deviation projection for melanoma spheroid, logarithmic scale). Shadow artifacts are indicated by the yellow dashed boxes in the OCM cross-section images (logarithmic scale). Scale bars: 200 µm.

The cross-section images (locations are indicated by the yellow dashed lines in the *en face* images) exhibit distinct shadow artifacts, as seen in the lower row of Fig. 3. For the zebrafish embryo, the eye with high melanin content and the yolk with high-attenuating internal structures cast shadows (signal loss) on deeper morphological features (marked by yellow dashed boxes in the OCM cross-section images). Similar artifacts were also identified in melanoma spheroids, where cells with high melanin levels limited the penetration depth due to high absorption. These shadow artifacts from a single viewing angle were also clearly revealed in the 3D rendering (see Supplementary Movie 4).

The *en face* images obtained by average intensity projection or standard deviation projection display the combined signal from different sample depths. Naturally, the artifacts caused by shadowing are not as obvious in this type of visualization. Fig. 4 demonstrates *en face* OCM images, with less noticeable shadowing, of a 5 dpf less-pigmented zebrafish embryo obtained from eight viewing angles. Whole-body *en face* data were obtained from three angles (indicated by the sub-image frame colors). Zebrafish features such as eye, otolith, yolk, muscle, notochord, and fins were discerned clearly by the OCM setup. Complementary zebrafish features were visualized from individual angles, but darker regions can be observed in the images, depending on the reorientation angles.

**Figure 4.**
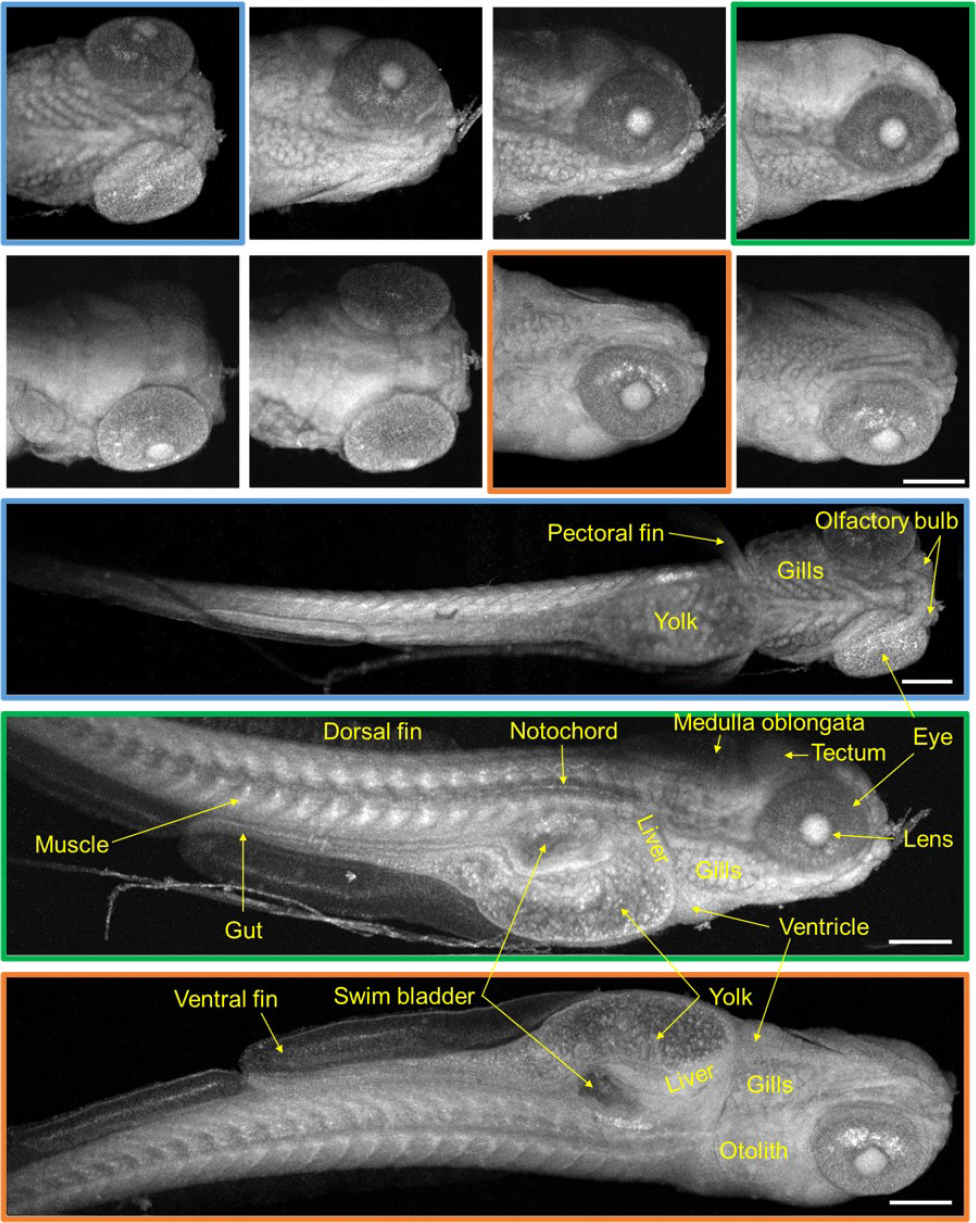
OCM *en face* images (average intensity projection, logarithmic scale) of acoustically reoriented 5 dpf zebrafish embryo (mutation *Mitfa*^*b*692*/b*692^/*ednrb1*^*b*140*/b*140^). Frame colors indicate corresponding angles. Scale bars: 200 µm.

### Reconstruction

The sample can be reoriented in a reproducible manner by acoustic actuation, but – in contrast to externally induced mechanical rotation of an object immobilized in a container – the exact orientation between the recorded OCM volumes is not known *a priori*. And, depending on the orientation, different parts of the sample are occluded due to attenuation by structures in the sample, which for a zebrafish embryo is especially pronounced for the eye and the yolk sac. Also, structures of different group-RI inside the sample cause a local delay or surge of the recorded A-scan [28, 35, 36]. This effect is especially visible as a delay for structures behind the lens portion of the eye.

Due to the distortion mentioned in the previous paragraph, the OCM volumes belonging to different orientation angles are not simply related by a rigid body transform, but correspond to each other in a more complicated manner. Moreover, the distortion and shadowing artifacts hinder a reliable registration of the orientations of the different volumes. Therefore, we formulate the fusion as an inverse problem, where the OCM image formation is expressed as a physical forward model. This approach grants us the flexibility to deal with these uncertainties by constraints and regularization. This includes total variation (TV) and Tikhonov regularization as well as positivity and object support constraints. We solve this inverse problem by means of a gradient-based optimization approach, whose optimization parameters consist in the underlying reflectivity map *R* as well as attenuation *α*, RI contrast Δ*n* and motion parameters *q* (rotation parameterized by unit quaternions) and *t* (translation). For a detailed description we refer to the Supplementary Methods.

Fig. 5 and 6 show the reconstruction results of ULTIMA-OCT from the head section of 3 dpf zebrafish-embryos, a wild-type and a less-pigmented mutation, respectively. In a comparison of the OCM volumes and the reconstruction of the reflectivity map *R*, both depicted in logarithmic scale, one clearly appreciates the benefit of the proposed approach. One can see that both specimens strongly attenuate the signal in the OCM volumes, whereas the reconstructed reflectivities do no longer show attenuation or distortion artefacts. In the zebrafish of the wild-type the scattering and absorptive structures contained within the total attenuation *α* are present across the whole head section, whereas for the mutated specimen the eyes account for most of the attenuation. Vertebrate eye-lenses exhibit a graded-index (GRIN) profile, which increases towards the center. As the the most prominent structures of RI map are those belonging to the eye-lenses we employ regularization that promotes smoothness, whereas for the attenuation and reflectivity map we use edge-preserving regularization. The values obtained for the RI using this method are consistent with values from the literature [37, 38]. The data-set of the wild-type zebrafish embryo consist of OCM recordings of 11 angles, which are roughly distributed between 0 and 360°, whereas for the less pigmented mutation zebrafish embryo 10 angles were used. 3D visualization of reconstructions comparing single- and multi-view of the wild-type zebrafish embryo can be found in Supplementary Movie 5 and 6 with a 3D rendered object and a flythrough-animation, respectively.

**Figure 5.**
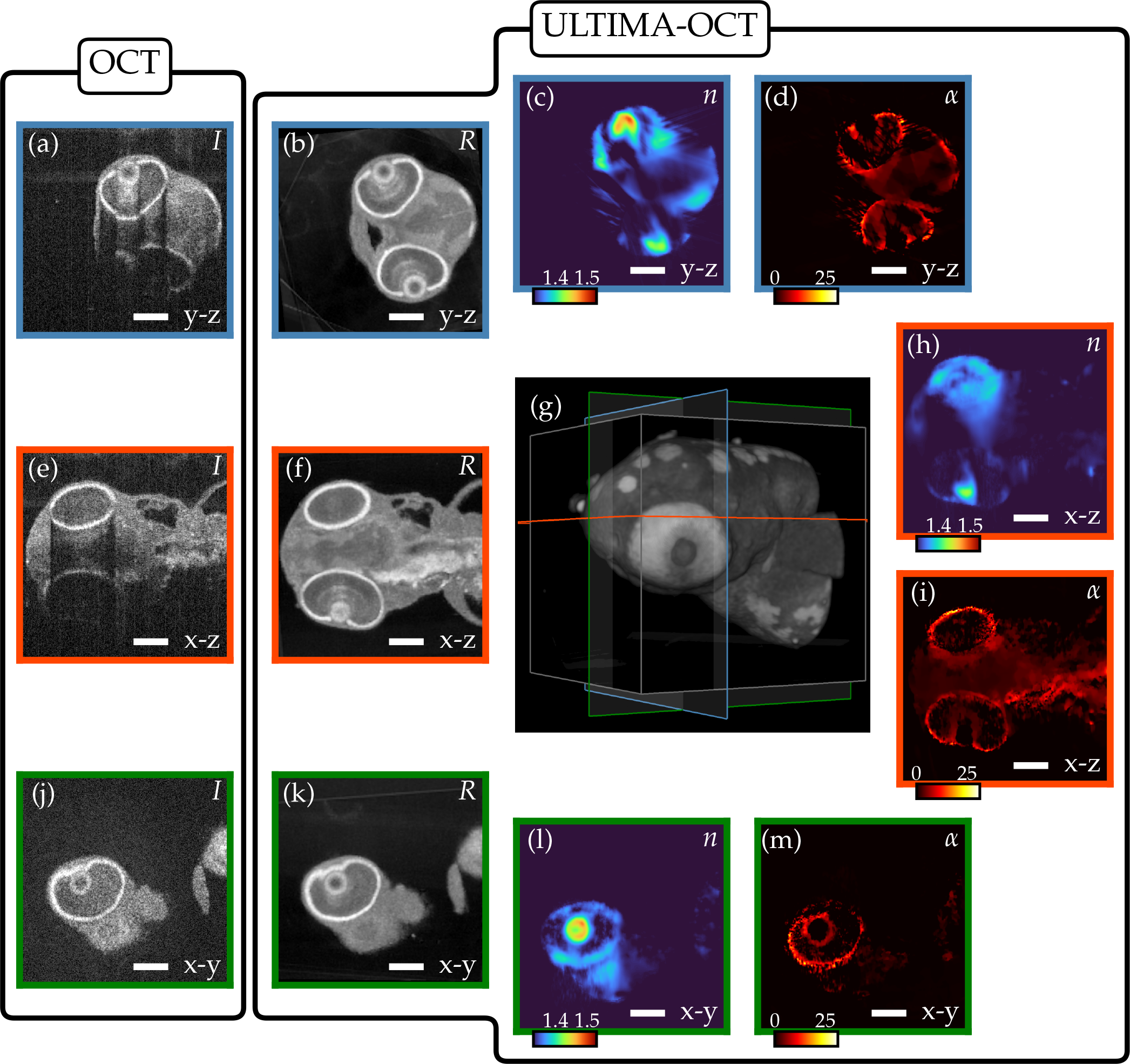
Reconstruction results for the head section of a 3 dpf wild type zebrafish embryo: In (a-d) the (*y z*), (e,f,h,i) the (*x z*) and (j-m) depict the (*x y*) sections of the reconstructions. The leftmost column (a,e,j) shows the sections of the recorded OCM volumes in logarithmic scale, whereas the adjacent column (b,f,k) show the reconstructed reflectivity map *R* of the same sections in logarithmic scale. (d,i,m) show the slices of the reconstructed attenuation map *α* (in mm^−1^), whereas (c,h,l) depict the sections through the reconstructed RI distribution *n*. In (g) a 3D rendering of the reconstructed reflectivity map is shown, together with the planes shown in (a-f,h-m). Scale bars: 100 µm.

**Figure 6.**
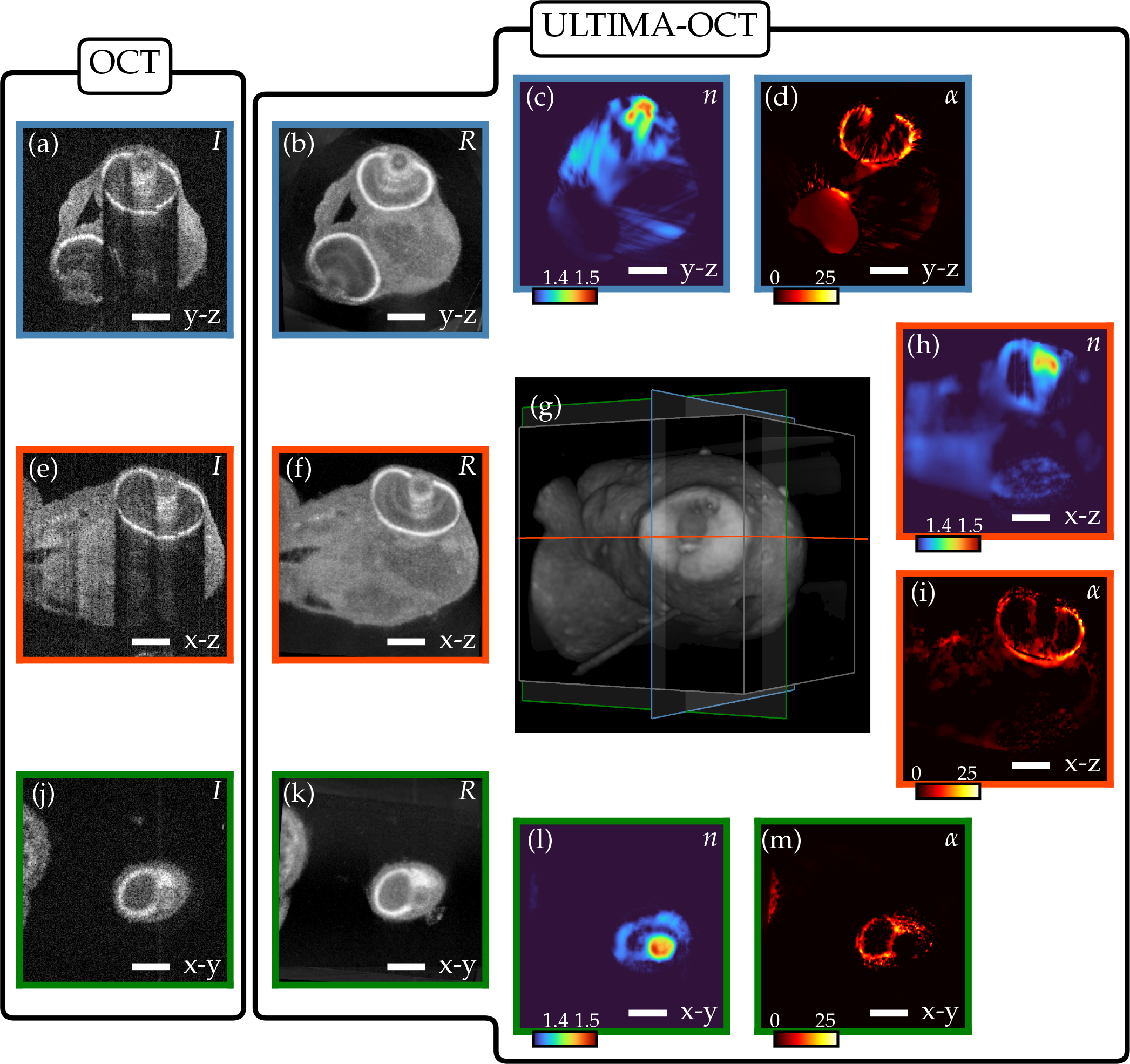
Reconstruction results for the head section of a 3 dpf zebrafish embryo of a mutation with reduced pigmentation: In (a-d) the (*y− z*), (e,f,h,i) the (*x− z*) and (j-m) depict the (*x− y*) sections of the reconstructions. The leftmost column (a,e,j) shows the sections of the recorded OCM volumes in logarithmic scale, whereas the adjacent column (b,f,k) show the reconstructed reflectivity map *R* of the same sections in logarithmic scale. (d,i,m) show the slices of the reconstructed attenuation map *α* (in mm^−1^), whereas (c,h,l) depict the sections through the reconstructed RI distribution *n*. In (g) a 3D rendering of the reconstructed reflectivity map is shown, together with the planes shown in (a-f,h-m). Scale bars: 100 µm.

## DISCUSSION

ULTIMA-OCT imaging and tomographic reconstruction provides volumetric information on the sample with enhanced penetration depth. This is achieved by acoustic reorientation, without the need for any moving mechanical parts, and in a non-contact way without the need of a supporting scaffold. This makes the sample much more accessible for mechanical probing and facilitates unobstructed and undistorted imaging. We will now discuss some current difficulties and limitations as well as possible extensions of the approach in the future.

The standard method to study samples such as live zebrafish embryos with OCT has been to manually position the embryo under anesthesia on a gel layer on a coverslip and image from one direction. The purpose of this gel layer is to lift the sample up to create sufficient distance from the coverslip, and the coverslip is usually tilted to get rid of the high reflection from the bottom coverslip. As a result, parts of the big sample may be out of focus and requires focus stacking or image stitching. ULTIMA-OCT has an acoustic chamber that can levitate samples up to 1 cm away from the bottom and top chamber surfaces, preventing these issues. Manipulating samples far from reflecting surfaces might not be possible for alternative acoustic strategies utilizing SAW as such devices are difficult to scale up. Further, we are unaware of stable reorientation of strongly asymmetric samples as zebrafish larvae using SAW, although demonstrated for the more axis-symmetric *C. elegans* sample [39].

While developing the approach we have used fixated specimens, but we expect no problems for live anaesthetized fish. Behavioral response under the influence of optical tweezers has successfully been studied in similar conditions [40]. OCT imaging of wake fish embryos does not seem feasible, since the considerably stronger acoustic forces necessary to stall an actively swimming fish will have non-negligible bio-effects (such as heating and cavitation), besides the problem of motion artifacts that will be inevitable. Importantly, in our platform during the acquisition times of 8 s to 23 s to capture a single angle volume of a field of view ranging from 0.67 mm*×* 0.67 mm to 0.76 mm *×*3.78 mm, the sample could be kept stable by acoustic trapping, i.e., we did not observe any motion artifacts.

The shape of the levitated specimen has an influence: For elongated and very asymmetrical samples such as the zebrafish embryos, we found that we can precisely control the orientation to basically any desired angle, as seen in Supplementary Movie 2. Reorientation leads to rotation around the major axis, but additionally there can also be a tilt due to asymmetry in the mass distribution and due to non-uniform acoustic forces in the chamber. This motion effect is dealt with by our reconstruction algorithm and does not impose a serious restriction. Since the targeted types of samples are never perfectly spherical, our manipulation strategy is also suitable for less asymmetric samples as shown in Supplementary Fig.4 and Movie 3, where we successfully manipulated melanoma spheroids. To be able to handle organoids in our platform, we would need the organoids in suspension or embedded in a gel droplet, which was not available to us at this stage. It has already been demonstrated that acoustofluidics can play a useful role in the trapping and merging of organoids [41], where organoids were formed in suspension or in matrigel followed by matrigel removal before acoustic manipulation and further growth in gel. In the future it may be possible to fully avoid the use of matrigel and to create protocols for organoid growth mediated by ultrasound, similar to growing cancer spheroids under acoustic trapping [42–44].

The range of trappable sample sizes depends on the design of the acoustic chamber. In the current manipulation chamber, we used a transducer resonance frequency corresponding to a wavelength in the water of 2.2 mm. In previous studies, we have found that we can manipulate samples up to a thickness slightly above *λ/*2 in our devices [9, 13]. We have demonstrated trapping of zebrafish up to 5 dpf (about 700 µm thickness), at a minimum voltage of the top transducer of 20 V corresponding to a maximum pressure amplitude of 80 kPa. For large samples relative to the trapping wavelength the acoustic radiation forces scaling with the sample radius gets more complicated [29, 31] than for the small particle limit [45]. However, we are not at the limits in driving voltage and we believe our platform is also suitable for handling larger samples up to a thickness of above 1 mm. Concerning the lower bound for trapping, we have demonstrated stable trapping and reorientation of a cancer spheroid of about 400 µm thickness, but the exact limits will be sample-dependent and need further investigation. To manipulate even smaller sample sizes at higher trap-stiffness, one could either use a higher harmonic frequency in the same device, or transducers of smaller size. Some more information on the choice of the size of the acoustic chamber can be found in the Supplementary Methods.

For long-term monitoring of live samples, it is necessary to implement a temperature control of the chip (e.g. by attaching a Peltier element), since the conversion to acoustic waves in the piezo-electric transducers leads to excess heat. In our current device, the relatively large aluminium plate sealing the back of the chamber is in contact with the fluid and transducers on one side, and can dissipate heat to the surrounding air. At large driving voltage, minimal changes in the acoustic resonances are observed, which indicate minor temperature changes. Further, for longer-term observations of developing cell spheroids, easy cell media exchange is important. It is straightforward to implement microfluidic channels to the front and the backside of the chamber. Biocompatibility has been verified for a multitude of similar acoustic trapping platforms [16, 46, 47], and our measured pressures are below the limits of which adverse bio-effects and cavitation are expected. However, we will need to assess the effects of our acoustic trapping for long-term experiments on the targeted live samples in more detail. Furthermore, biological samples are pushed to low-pressure regions, and here we often operated the transducers at higher voltages than strictly necessary for levitation and reorientation. Thus, with fine-tuning of the acoustic parameters, the chamber-dimensions and the acoustic resonances, it should be possible to operate at even lower pressures. Using different transducers with a slightly higher trapping frequency could also be possible, which further limits the likelihood of cavitation.

The targeted samples are addressable by OCM, with the penetration depth depending on the light source’s wavelength and the sample’s scattering and attenuation properties. Longer wavelengths allow for deeper penetration at the cost of lower resolution. High-scattering and high-attenuation structures can obstruct the incident beam, resulting in shallower penetration. Therefore, OCT imaging depth is limited to 1 mm to 2 mm for most biological samples, and OCM can even be less. To achieve high lateral resolution, the numerical aperture is increased in OCM. However, this can result in a decrease in the depth of field and lead to non-uniform lateral resolution at varying depths. Ultrasound imaging can reach deeper but has a poorer resolution. ULTIMA-OCT maintains the high resolution of OCT/OCM and compensates for shadow artifacts, thereby reconstructing the sample part beyond the limits of a single angle’s penetration.

The reconstruction in multi-angle OCT previously has been performed on static samples [28] or with known motion [36] using a number of angles that is an order of magnitude larger than in this work. Furthermore, the uncertainty in the orientation of the sample adds additional ambiguity, which makes the reconstruction process even more challenging. To achieve sufficient accuracy on the registered angles we made use of compressive algorithms in the reconstruction process. In the first stage of the algorithm, it is crucial to make heavy use of regularization on the reflectivity map *R* to deal with the unknown motion in-between volume recordings. To make this step efficient, coarser representations of the recorded data can be used to obtain a first guess of the reflectivity, RI, attenuation map and motion. After the motion is registered with sufficient accuracy, the high resolution reconstruction can be initialized with the parameters estimated from the first stage. Additionally, the strength of the regularization (explained in the Methods Section and in more detail in the Supplementary Methods) can be lifted in order to also record fine-grained structures and to make use of the full resolution in the OCT data-set.

Limitations of the presented approach concern the attenuation and RI maps. In modeling the attenuation we assumed the scattering and absorption to be independent of the recording angle. While the attenuation map is a useful quantity for the fidelity of our model, the angular independence might not be given and therefore the reconstruction of *α* is limited in its informative value. As the number of orientations used in this work is small, the reliability of the RI-map outside the eye regions may be restricted. Although the RI values of the lens portion of the eye can be estimated accurately, the reconstruction of the RI map is strongly dependent on the available angular coverage in the data-set and the specimen itself. As a change in the RI at a location manifests itself as a delay in the structures behind that location, distinct structures have to be visible in the OCT signal.

ULTIMA-OCT combines cutting edge modalities in acoustic actuation, OCT and model-based tomographic reconstruction. Our 3D printed low cost chamber is a simple add-on to an OCT imaging platform permitting multi-angle acquisition of a variety of samples in a non-contact manner. Since the reconstructed maps for reflectivity, attenuation and RI are tightly linked physically, future work on the image fusion could entail utilizing a more refined physical forward model in the form of a unified treatment of those quantities. Our strategy of multi-angle OCT could potentially also profit from a full wave-optical treatment of the light-matter interaction [48]. The presented strategy of step-wise reorientation, registration and reconstruction can be applied to a wide range of imaging modalities in microscopy. Thus, we believe that this work has great potential for in *in-vivo* imaging in biomedical research.

## METHODS

### Acoustic manipulation chamber

The chamber frame with a symmetric octagon cross-section is 3D-printed (Original Prusa i3 MK3, Prusa Research, Czech Republic) in a polymer (PET-G, RS: 891-9309, Austria) with open front- and backside and with windows around the 8 sides for attaching 4 piezo-electric transducers and 4 reflectors. The 4 transducers are positioned on the top part of the chamber, as shown in Fig. 2b and Supplementary Fig. 1 (top-transducer and S1-S3 side-transducers) with the 4 reflectors on the opposite parallel side. For imaging compatibility through the bottom of the chamber, a 170 µm thick coverslip seals the bottom and acts as the reflector of the acoustic waves from the top-transducer. The remaining three reflectors (R1-R3 in Supplementary Fig.1) are machined in aluminium or cut from a 170 µm coverslip. All 4 transducers are (8 mm *×* 15 mm, 3 mm thick) plate transducers made of Lead Zirconate Titanate (PZT) (Pz26, CTS Ferroperm, Denmark). With the aim of levitating samples of a size in the mm-range, we chose these transducers with a thickness resonant mode frequency of the bare transducer around 670 kHz and a wavelength in water of about 2.2 mm. Resonantly enhanced bulk acoustic waves generate standing waves of sufficient forces in our chamber to levitate and reorient our samples. The specific chamber height used here (19.2 mm) was found by an iterative approach of adjusting the chamber dimensions of the 3D printed frame based on simulations and characterizing the acoustic resonances by electrical impedance measurements and experiments, see Supplementary Fig. 2 for specific chamber dimensions and detailed information in Supplementary Methods.

The backside of the printed chamber frame is glued to a machined octagon-shaped aluminium plate to seal the back of the chamber. To attach this aluminium plate, transducers and reflectors to the printed chamber frame, and make the chamber water-tight we use standard (cyanoacrylate) super-glue followed by nail-polish. The bottom silver plated electrode of each piezo-plate is connected to the aluminium with silver paint (RS: 123-9911, Austria) for thermal and electrical connection (common ground). To electrically connect to the top-and the bottom electrodes on each transducer we use copper wires and silver-paint to the top electrode and aluminium plate respectively (Supplementary Fig. 1). We drive each transducer with an AC signal from waveform generators: two single output waveform generators (Agilent 33220A, RS components, Austria) to drive side-transducers S1 and S2, and one dual output waveform generator (Keysight 33522B, RS Components, Austria) to drive the top- and S3-transducer. Each signal is amplified by power amplifiers (eval-ADA4870, RS: 836-8714, Austria) and impedance matching transformers, see details in Supplementary Methods. To ensure levitation of our samples we operate the top-transducer above 20 V, and to reorient the samples we tune the voltages of the transducers in the range of 20 V to 35 V. This corresponds to maximum pressure amplitudes in the range of 80 kPa to 150 kPa measured in the anti-node with a hydrophone (NH0200, Precision Acoustics, United Kingdom), see Supplementary Methods. Please note that voltage refers to peak-to-peak voltage throughout the paper.

### Operation for multi-angle imaging

The assembled octagon chamber is placed in a 3D-printed sample holder (Supplementary Fig. 1) that fits on the inverted microscope stage, attached via an adapter above the objective in the imaging setup in the case of OCT. We tilt the holder 90° so the open chamber front faces upwards and rinse the chamber with a 0.1 % Triton X-100 solution (Merk, Germany) to make the chamber more hydrophilic to limit bubble formation at surfaces when filling the chamber. We then fill the chamber with the liquid (distilled water, tap water or 1X PBS), place the sample inside and seal the front with a coverslip. This front coverslip is kept in place by adhesion forces, and can easily be removed for sample exchange. The top transducer is turned on and we tilt the chamber to levitate the sample in the middle region of the chamber. The holder is placed on the imaging stage, and we start the acoustic manipulation to reorient the sample and acquire images through the bottom coverslip of the chamber at each desired step. Dark-field image acquisition (see Supplementary Methods for setup details) was used to optimize the acoustic settings for stable reorientation, before performing OCM. The top transducer is “far away” from the trapped sample (about 7 mm to 12 mm), but to further limit the back-reflection from the top transducer during OCM, we color the bottom silver plated electrode with black ink, which does not affect its acoustic performance.

### Samples

#### Zebrafish (*Danio rerio*) embryo preparation

In this study we used zebrafish of the pigmented wild-type Tubingen strain (wild-type) and double mutant transparent *Mitfa*^*b*692*/b*692^/*ednrb1*^*b*140*/b*140^ fish with reduced pigmentation. After spawning, eggs were maintained in egg water at 28 °C under standard conditions for up to 3 and 5 dpf. After overnight fixation in 4 % paraformaldehyde (PFA), embryos were washed with phosphate-buffered saline (PBS) and were stored at 4 °C until they were used for imaging.

#### Spheroid preparation

The murine melanoma cell line B16-F10 was used to form spheroids by the hanging drop method. Cells were grown in Dulbecco’s Modified Eagle Medium supplemented with 10 % fetal bovine serum, 1 % penicillin/streptomycin in a humidified incubator at 37 °C, 5 % CO_2_. When 80 % cell confluency was reached, cells were trypsinized, and 1000 cells per 25 µl media were placed as droplets onto a Petri dish lid, inverted, and returned as top of the Petri dish bottom part, which was filled with 15 ml PBS for humidity and incubated for 4 days. During this time, individual spheroids formed in each droplet. The spheroids were collected in microcentrifuge tubes, washed 3 times with PBS between centrifugation (300*×* g, 5 min), and fixed with 4 % PFA at room temperature for 20 min before washing 3 times again with PBS. Spheroids were then stored at 4 °C until use.

### Adaptation of OCT set-up

The setup [49] (as illustrated in Fig. 7) used in this work is a spectral domain OCM system using a compact polarization-aligned three-superluminescent-diode laser source (EBD290002, EXALOS AG). The OCM laser source had a center wavelength of 845 nm and a wide bandwidth of 131 nm, resulting in a high axial resolution of approximately 3.7 µm in air, corresponding to 2.68 µm in tissue (with a RI of 1.38). The lateral resolution of the OCM system was about 3.4 µm. Through reflective collimator 1, the OCM beam was directed to the OCM system and then divided by a beam splitter to the sample arm and reference arm, respectively. The laser power on the sample was around 1.53 mW. Reflective collimator 2 was connected with a homemade spectrometer [50, 51] to capture the interferogram of backscattered light from the reference mirror and the sample. The acoustic chamber was mounted on a chamber holder and implemented in the OCM system using a 3D-printed adapter. Precise sample positioning and focus adjustment were achieved using a three-axis translation stage (MAX313D/M, Thorlabs).

**Figure 7.**
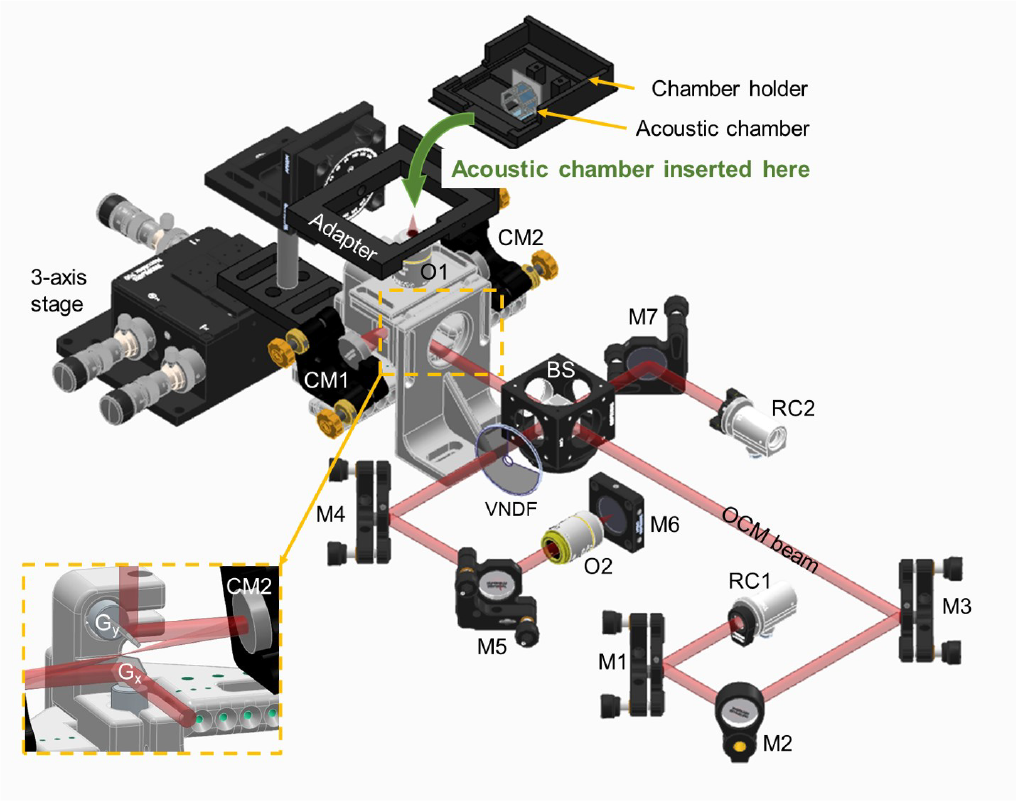
The schematic of the OCM system and the add-on acoustic chamber. RC: reflective collimator; M: mirror; BS: beam splitter; VNDF: variable neutral density filter; CM: concave mirror; O: objective; *G*_*x*_: x galvanometer scanner; *G*_*y*_ : y galvanometer scanner.

Volumetric OCM data was obtained by the raster scanning of a pair of galvanometer scanners. During imaging, a 20 kHz camera line scan rate of the OCM spectrometer was used, corresponding to a sensitivity of 107 dB. For zebrafish embryo imaging, a scanning step size of 1.68 µm was used for smaller field-of-view imaging (head region), and 2.52 µm was used for whole fish imaging. A scanning step size of 1.68 µm was employed for melanoma spheroid imaging.

After standard OCT preprocessing steps (background subtraction, resampling, digital dispersion compensation, fast Fourier transform, and logarithmic calculation), OCM raw binary data was converted to three-dimensional images [52]. *En face* images of the zebrafish embryos were obtained by average intensity projection, and the *en face* image of the melanoma spheroid was obtained by standard deviation projection using Fiji [53]. To create the OCM cross-section image of the zebrafish embryo, 7 B-scans were averaged consecutively. Similarly, the OCM cross-section image of the melanin spheroid was obtained by averaging 3 consecutive B-scans. 3D rendering of the volumetric data was achieved using Amira 3D (Thermo Fisher Scientific, version 2023.1.1).

### Reconstruction algorithm

The data-set consisting of OCM volumes of the specimen at multiple different orientations serves as the starting point for the reconstruction algorithm. We follow a model-based approach to describe the observed OCM data by the interaction of a reflectivity *R*, attenuation *α* and RI contrast map Δ*n*. Inspired by the works of [28, 36, 54], the detected OCM signal *I* is modeled line-wise by a layer-by-layer based propagation

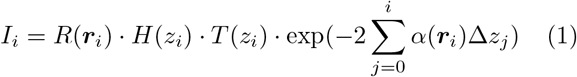

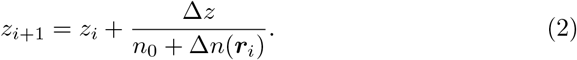

Starting with ***r***_0_ as the boundary conditions for the coordinates, the signal is traced through the specimen represented by *R, α* and Δ*n. T* and *H* denote the confocal point spread function and sensitivity roll-off, respectively [55].

To extract the maps for *R, α* and Δ*n* we formulate the reconstruction as an optimization problem. The error metric we aim to minimize is composed of data fidelity, total variation, and *l*2-norm, as well as positivity constraints on *R, α* and *n*. In addition to *R, α* and *n*, also ***q*** and ***t***, rotation parameterised by quaternions and translations, represent optimization parameters. We solve the optimization problem jointly with Stochastic Gradient Descent (SGD), where we first extract motion parameters and reconstructions on a low-resolution representation. Afterwards, the high-resolution reconstruction is initialized with the resulting obtained low-resolution quantities, and further refined iteratively to yield the final reconstruction. For a detailed explanation, we refer to the Supplementary Methods.

The numerical optimization was conducted in JAX[56] 0.4.13 using Python 3.11 on a workstation equipped with an Nvidia RTX 4090 GPU. To obtain the low-resolution reconstructions we ran the algorithm for 2500 iterations and use a 3x smaller resolution (134^3^ and 150^3^), whereas for the high-resolution reconstructions (400^3^ and 450^3^) 250 iterations were performed. An iteration represents the a single pass through the whole dataset. The whole reconstruction process of took approximately 20 min.

## FUNDING

This work was supported by the Austrian Science Fund (FWF) under SFB-grant F68 *Tomography across the scales* (sub-projects F6803-N36 and F6806-N36), H2020-ICT-2018-20 project REAP with grant agreement ID 101016964, and the Joint Ph.D. Program Medical University of Vienna/NTU Singapore “Kooperation Singapur” (Grant No. SO10300010).

## ACKNOWLEDGMENTS

We thank Nicole Schmitner (Institute of Molecular Biology, University of Innsbruck) for providing us with zebrafish embryos and describing the features, as well as for valuable discussions, Abigail J. Deloria (Center for Medical Physics and Biomedical Engineering, Medical University of Vienna) for providing us with melanoma spheroids.

## DATA AVAILABILITY STATEMENT

To be specified.

## SUPPLEMENTARY INFORMATION

The Supplementary Material for this article can be found online at: to be specified. It covers details on acoustic chamber design and assembly, as well as an example of a reoriented melanoma spheroid. Moreover, further details on the reconstruction algorithm are included, and a list of supplementary figures and movies.

## AUTHOR CONTRIBUTIONS

WD, MRM, and SM conceived the general idea of the work. MKL developed the acoustic manipulation strategy, designed and fabricated the acoustic chamber. MKL and SD carried out the acoustic-OCT experiments at the Medical University of Vienna. SD, RL, and WD planned the OCM experiments, SM and RL developed the numerical reconstruction algorithm, and all authors contributed to the structuring and writing of the paper. rotation of biological samples in a sono-optical microfluidic device, Lab on a Chip **21**, 1563 (2021).

## Notes

### Competing Interest Statement

The authors have declared no competing interest.

