## Supplementary Material for "Ultrasound-Induced Reorientation for Multi-Angle Optical Coherence Tomography"

(Dated: September 18, 2023)

Here we present supplementary figures, methods and movies that we provide to support the main article. The document covers details on acoustic chamber assembly, chamber dimensions and acoustic resonance characterization. We also describe the results of the acoustic pressure measurements with a needle hydrophone and the details of the setup for darkfield imaging. And we include more information on the acoustic actuation and the underlying mechanisms (acoustic torques), and give an additional example, the reorientation of a melanoma spheroid. Moreover, further details on the reconstruction algorithm can be found.

**Contents:**

**Supplementary Figures**

**Supplementary Methods**

**Supplementary Movie list**

**Supplementary References**

* contributed equally

### I. SUPPLEMENTARY FIGURES

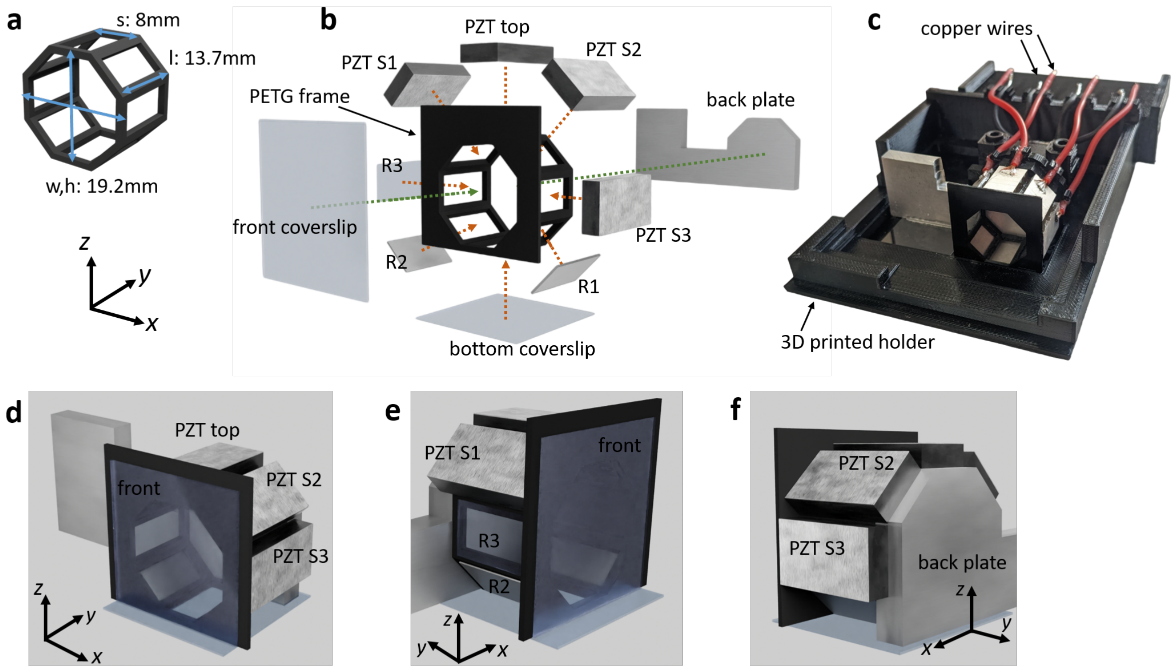


**Supplemenatry Figure** 1. **Dimensions and chip assembly: a** The 3D printed chamber frame has a symmetric octagon cross-section with a height of 19.2 mm ($z$-axis), sides of 8 mm and a length of 13.7 mm ($y$-axis). **b** Illustration of assembly of the chamber: 4 piezo-electric plate transducers and 4 reflectors are glued around the 8 sides of the octagon frame. The front and back is sealed with a coverslip and an aluminum plate, respectively. **c** Image of the assembled chip in a 3D printed frame compatible with the microscope stage. **d-f** Illustrations of the assembled chip from three viewing directions.

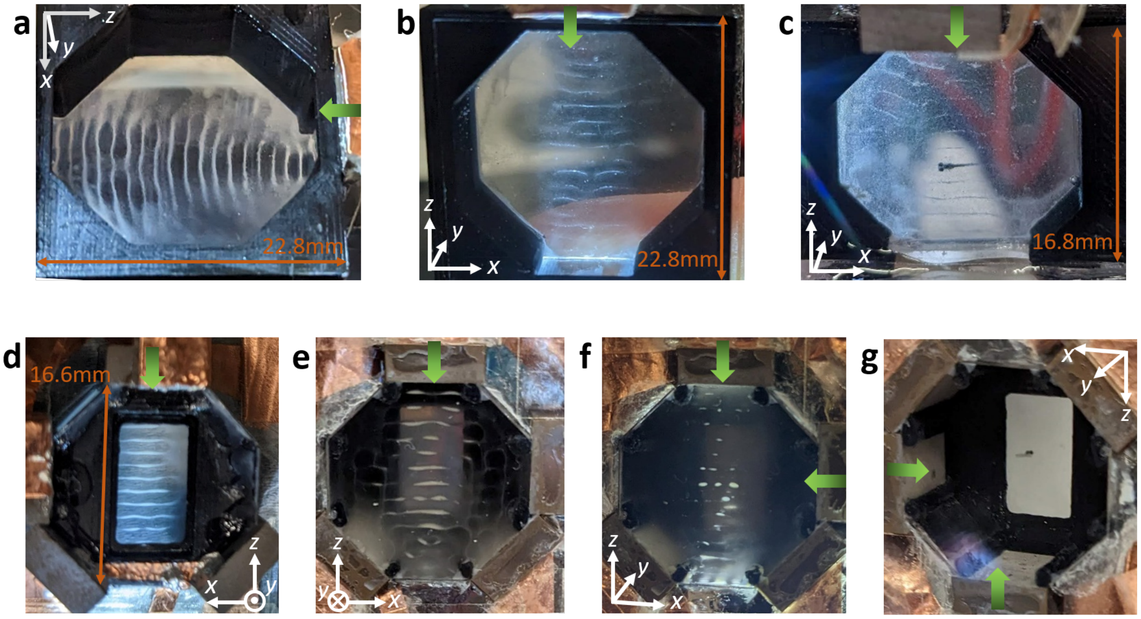


**Supplemenatry Figure** 2. **Mode visualization and performance: a-f** Acoustic trapping of a yeast cell suspension to visualize mode-patterns at measured resonance frequencies. **c** A zebrafish embryo (3 dpf) is in addition levitated and **g** only a 3 dpf zebrafish is trapped. The indicated $x$,$y$,$z$-axis are with respect to the chamber as in Supplementary Fig.1, where the $z$-axis is parallel to the height of the chamber and the standing wave from the top-transducer, $x$-axis to the width of the chamber and the standing wave from the side-transducer (S3). The acoustic propagation direction of active transducers is indicated with green arrows. We manufactured a range of octagon sizes initially to optimize the dimensions for resonances in the fluid of sufficient forces to levitate large samples such as zebrafish embryos with the top transducer only. The octagon frame in **a** and **b** has a height of 22.8 mm, **c** of 16.8 , and the octagon frame shown in **d-g** has a height of 16.6 mm. **a-e** Only the top transducer is active. The acoustic standing wave (parallel to $z$-axis) is perpendicular to the direction of gravity in images in **a** and **d**, and in the other images, the standing wave is parallel to direction of gravity. **f** and **g** Trapping with both top transducer and orthogonal side-transducer (S3) active. Hence the yeast cells on the coverslip are further confined from stripes in **e** to islands in **f** (from planes to cylinders in 3D). And further in **g**, the zebrafish is oriented with its long axis to the length of the octagon, compared to the top-transducer only trapping in **c**.

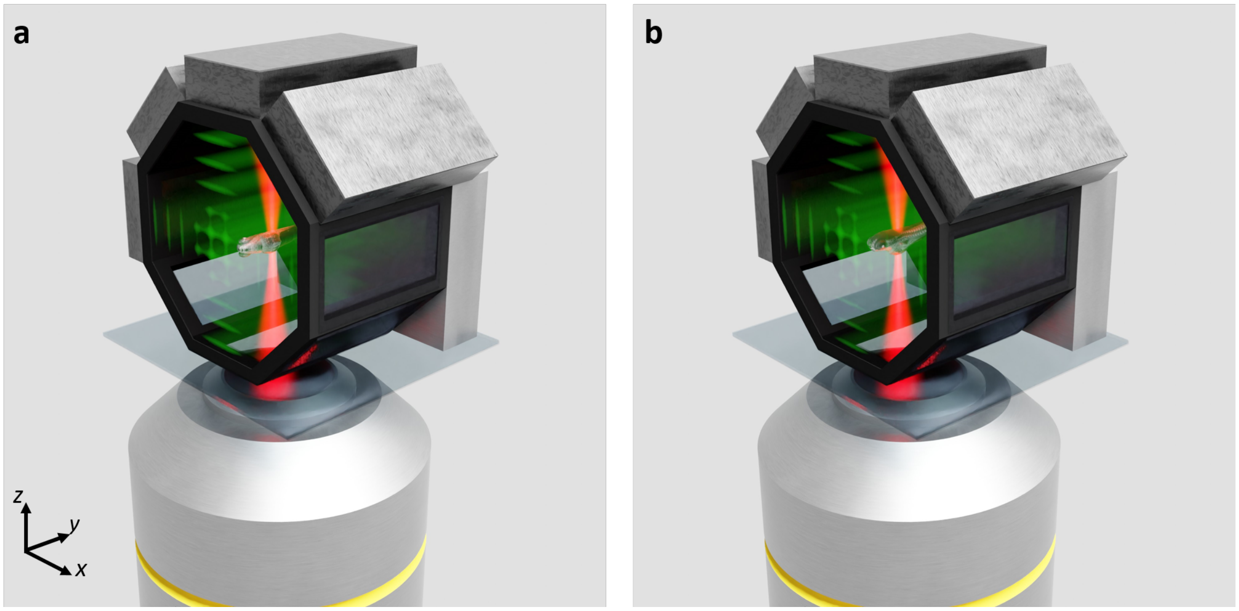


**Supplemenatry Figure** 3. **Working principle of ULTIMA-OCT:** Acoustic manipulation actuated by a 4- or 2-transducer approach leads to stable reorientation of the sample 360° around its major axis, at a step-size determined by the adjusted step-size in amplitude and/or phase-shift between the active transducers. Acoustic waves are illustrated in green and the imaging beam in red. **a** With a dominant top-transducer, the zebrafish embryo is oriented with its minor axis to the steepest trap-stiffness in the $z$-direction with one eye down and the yolk to the left or the right. **b** With a dominant S3 transducer, the zebrafish is aligned with its minor axis to the steepest trap-stiffness in the $x$-direction typically with its heavy yolk down due to gravity. By adjusting both the phase-shift and the voltage between the two orthogonal transducers, we can reorient the sample to any desired orientation, and at each desired angle we scan the OCT beam to record multi-angle 3D imaging data as illustrated in the animation in Supplementary Movie 1 and in the recording in Supplementary Movie 2. Note that this figure and animation is not to scale and does not fully resemble the actual assembly of the used device in Supplementary Fig.1.

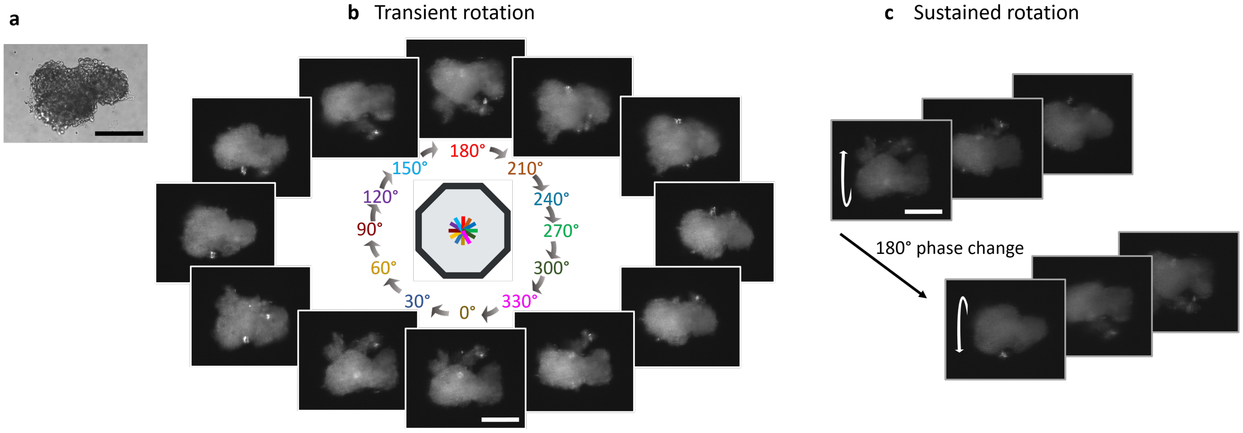


**Supplemenatry Figure** 4. **Reorientation and sustained rotation of melanoma spheroid:** **a** Brightfield image of the melanoma spheroid (on coverslip). **b** Darkfield images of the melanoma spheroid which is acoustically reoriented to stable trapping orientations around its major axis in the octagon chamber. Here 12 different orientations are shown, but a finer angle step-size is possible. The indicated angles are approximate. **c** We can also induce a sustained rotation of this sample and by changing the relative phase by 180° between the two transducers generating the spinning torque, we can flip the rotation direction (see Supplementary Movie 3). Scale bars: 250 µm.

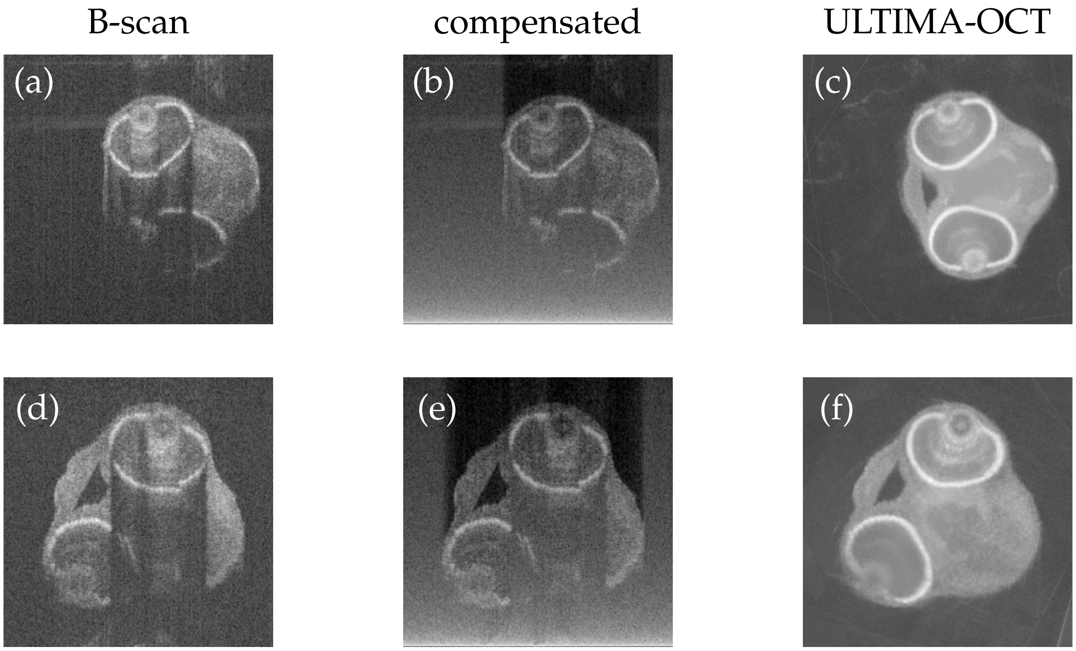


**Supplemenatry Figure** 5. Comparison of ULTIMA-OCT with an established shadow compensation algorithm. (a,d) show the raw B-scans from the 3dpf wild-type and mutated, respectively. (b,e) show the results applying the shadow compensation from [1], while (c,f) show the reconstruction from the proposed method.

#### II. SUPPLEMENTARY METHODS

##### A. Assembly of octagon chamber

The (8 mm × 15 mm, 3 mm thick) transducers (PZT top and PZT S1-S3 in Fig.1) are attached with the long side aligned with the length of the printed octagon frame ($y$-axis), covering the open frames around the sides. The backside of the octagon chamber is sealed with a hand-milled octagon shaped back-plate. The reflector for top-transducer is the bottom coverslip in Supplementary Fig.1**b** which is a standard 170 µm coverslip for imaging compatibility from the bottom. The reflector R3 is cut from a standard 170 µm coverslip, and the reflector R1 and R2 are machined in aluminium. All transducers, reflectors and back-plate are glued to the octagon frame with standard (cyanoacrylate) super-glue. After the glue has dried, we also add a layer of standard nail-polish around the edges of the components interfacing the octagon frame to ensure that the chamber is water-tight. The transducers extend $\approx$ 1.5 mm over this back-plate, and we add silver-paint between the bottom electrode and the aluminium plate for thermal and electrical connection. With this the back-plate acts as an electrical common ground for the transducers bottom electrodes, and also to conduct heat away from the chamber as the inefficient piezo-electric conversion can lead to excess heat. The back-plate is extended in size in the $x$-direction and implementation of a Peltier element for more precise temperature control for studying live samples in the future would be straight forward. Electrical connection to the transducer electrodes is made via copper-wires to the top electrode and the back plate, respectively, with super-glue and silver-paint, seen in the image of the assembled device in Supplementary Fig.1**c**. The front coverslip is placed covering the front of the chamber only after filling the chamber with liquid and sample, and hence kept in place by the adhesion forces for the duration of the experiments. This is an easy approach for this proof-of-principle, and micro-fluidic in-lets and out-lets are planned modifications for future devices.

##### B. Chamber dimensions and characterization

Due to the open frames around the sides of the octagon frame the transducers are in direct contact with the fluid during experiments. We operate the transducers close to the first harmonic thickness resonant mode frequency of the bare transducer around 670 kHz.

Further, to achieve sufficient forces to levitate heavy samples such as 3-5 dpf zebrafish embryos, we rely on resonant enhancement of the bulk acoustic waves in the fluid cavity. We perform simulations based on a simple one-dimensional model [2] to choose the initial chamber dimensions. To experimentally determine the resonances in our chamber, we perform electrical impedance measurements of the transducer while scanning the frequency with a Red Pitaya V1.1 device (RS components, Austria) with power amplifier (EVAL-ADA4870), see [3] for more details. By comparing the electrical impedance scans of each transducer between an empty and a filled chamber, we get a good indication of the strongest resonances in the fluid-filled chamber. The specific octagon height used to collect data in this study (19.2 mm) is found by printing various sizes of octagon frames based on the simulations, and on the results from the electrical impedance measurements and experiments verifying whether the forces from the top transducer are sufficient to levitate the sample, in addition to visualize the mode-pattern by trapping a yeast cell suspension, as seen in Supplementary Fig.2. In **a** the acoustic standing wave is perpendicular to gravity and the accumulation of yeast cells on the bottom coverslip gives a better contrast. In the other sub-images (**b-g**), with the acoustic standing wave parallel to direction of gravity, it is more difficult to levitate small samples as yeast cells with the chosen wavelength, and only accumulations of cells can be levitated close to the cover-glass due to secondary acoustic radiation forces and friction forces. Hence, this visualization with yeast cells is not fully 3D. The stripy mode-pattern seen with trapping by only the top-transducer in Supplementary Fig.2**a-e** is further confined along the $x$-axis when the orthogonal S3 transducer is active in addition (corresponding to a cylindrical trapping pattern in 3D). The acoustic radiation forces scale with the volume of the trapped particle when the sample radius is much smaller than the trapping wavelength [4], and is more complicated closer to this limit [5, 6]. The zebrafish sample can be levitated with the top transducer only in Supplementary Fig.2**c**, where the its major axis is typically aligned with the $x$-axis. With orthogonal S3 transducer active in addition (in sub-image **g**), the zebrafish major axis is aligned to the $y$-axis.

3D printing of the chamber frame therefore is a low cost and efficient way of producing a range of chambers to fine-tune the acoustic response, at a sufficient accuracy. As a measure of the printing accuracy of the symmetric octagon frame: the measured transducer resonances in the fluid are the same for the two pairs of transducers with the same glass and aluminium reflector, respectively, see II E for more details. The 8 mm width of the transducers sets a lower limit for the size of the printed octagon frame to fit our chosen transducers around the sides. Further, to efficiently excite the thickness mode of the transducer, a more plate-like geometry of the transducers are preferable, hence reducing the transducer size much further has some downsides. For our targeted samples, it could also be possible to use higher trapping frequencies around 1 MHz, for instance going to 2 mm thickness of smaller transducers, and reducing the octagon frame size correspondingly.

##### C. Signal generation and acoustic pressure measurements

The AC output from the signal generators are each connected to a power amplifier, and we use an impedance matching transformer between the amplifier's 10Ω impedance output and each transducer to amplify the signal and match the transducer and amplifier impedance for increased power transfer. To initially levitate the samples with the top transducer, we operate the transducer in range of 30 V to 35 V and the sample stays levitated at voltages down to 20 V. We note that all driving voltages refer to peak to peak voltages measured across the transducer. To characterize the resulting pressures in our chamber we carried out hydrophone measurements with a needle hydrophone (NH0200, Precision Acoustics, United Kingdom). The chamber was tilted 90° and filled with distilled water, and the needle hydrophone was attached to a 3D stage and positioned into the center of the open front of the chamber. We localized the maximum pressure amplitude by scanning in the acoustic propagation direction and performed measurements at multiple voltage settings of the top transducer. Our typical operating voltages between 20 V to 35 V result in a maximum pressure amplitude between 80 kPa ± 10 kPa and 150 kPa ± 16 kPa where the uncertainty is the standard deviation including measurement uncertainty and the 9 % uncertainty of the hydrophone measurement stated by the manufacturer. The underestimation in the measured relative amplitude due to hydrophone directional response for 1 MHz (stated by manufacturer) is taken into account, and directional response for 0.6 MHz is expected to be less. Notably, these pressures are well below the limits of recommended Mechanical Index (MI) to limit the likelihood of inertial cavitation, and below the ranges reported in other studies where no adverse bio-effects have been reported [7]. The pressures reported

here are measured in the pressure anti-nodes of the standing wave which corresponds to the maximum pressure and pressure gradient associated with higher likelihood of cavitation and bio-effects [7]. The sample is trapped in the pressure nodes which creates a distance between the sample and potential oscillating bubbles and associated streaming effects.
In this study, we first used partially degassed distilled water (degassed by heating followed by cooling in an closed container) to limit bubbles at the non-smooth edges of the printed octagon frame when initially filling the chamber. As small bubbles disrupt the trapping, we have a clear indication that we have a bubble-free chamber. To demonstrate trapping in media with a higher gas-content, we needed to coat the inside of the chamber frame and between the printed frame and components (with nail-polish followed by the Triton rinsing) to avoid bubbles when initially filling the chamber. Starting with a filled, bubble-free chamber, the stable trapping was performed in tap water and 1X PBS buffer. As a precaution to avoid bubble formation due to the lower gas-solubility of liquid at higher temperature, we heat the liquid to $5^{\circ}$ more than room temperature before filling. A stable temperature control of the liquid and chamber in the future would alleviate this issue. Hence, we do not expect trapping in gas-perfused media to be an issue for our settings and future devices will be made with smoother surfaces and less edges to limit that gas is being trapped and more suitable hydrophilic materials or coatings could be used. Further, in this proof of principle we often operated the transducers at a higher voltage than strictly necessary, which can be avoided by fine-tuning the parameters if necessary. Using higher trapping frequency (around 1 MHz for instance) will also reduce the likelihood of cavitation effects.

##### D. Darkfield imaging

The optimization of the acoustic actuation was carried out on an inverted microscope (Zeiss AxioVert 200M, Zeiss, Germany) with optical image acquisition using a 5× objective (Zeiss Fluar 5×, NA 0.25, Zeiss, Germany), with sufficient working distance to image the center region of the chamber. We captured darkfield images with oblique illumination with LED (M625L4 - 625 nm, Thorlabs Inc, Germany) through the front coverslip of the chamber. The camera employed is a MatrixVision mvBlueFox3-2071a with a Sony IMX428 sensor (Matrix Vision GmbH, Germany). To capture the whole body of the zebrafish, as presented in the main article, we acquire two images at different $x$-positions of the stage that are later stitched together with Fiji [8].

##### E. Acoustic actuation

For the top and S3 transducer we use a dual output signal generator which allows us to drive the two transducers at the same frequency, but with different amplitude and a set relative phase. The two remaining side-transducers are each driven with a single output waveform generator. Because the octagon is made symmetric in this device, all transducers have a similar resonance frequency in the fluid-filled chamber. The top and S3 transducers have the same (coverslip) reflector, the S1 and S2 transducers have the same (aluminium) reflector, and the measured resonance frequency in the fluid-filled chamber is the same for the two pairs of orthogonal transducers (600 kHz and 590 kHz respectively). In each of the acoustic standing waves from the transducers there is a pressure node every $\lambda/2$, hence 15 nodes in each of the four directions. To manipulate the sample with all transducers, we levitate the sample in one of the five nodes in the vertical direction in the center region of the chamber, where the standing waves from all the transducers intersect. To do this we tilt the chamber filled with liquid and a sample with only the top transducer active, until the sample levitates in the center region. In future devices, we plan to implement micro-fluidic inlets and outlets to make loading and unloading of a new sample more efficient. When we then increase the amplitude of one side transducer, we orient the samples major axis to the length of the chamber. In this position, we now tune the relative amplitudes between the transducers, hence the magnitude of the acoustic radiation forces in each direction to rotate the sample.

The acoustic radiation forces acting on an asymmetric sample can lead to acoustic torques acting on the sample. We split the acoustic torque into two torque contribution: the acoustic restoring torque and the acoustic spinning torque [9]. While the restoring torque will align an asymmetric samples minor axis to the steepest trap-stiffness, the spinning torque, arising from acoustic absorption in the viscous media and in the sample itself can lead to sustained rotations of samples when the spinning torque is larger than the restoring torque. To generate a spinning torque, two orthogonal modes of the cavity have to be excited at the exact same frequency. Further, the spinning torque depends on the phase-shift between the two orthogonal transducers driving it [10]. To induce transient rotations for OCT imaging, and not sustained rotation, we operate at settings where the magnitude of the restoring torque is always larger than the spinning torque [9, 10]. We designed the octagon chamber to reorient the sample using the 4 standing waves, but also explored another strategy using only the the top transducer and the orthogonal S3 transducer. Using the four transducer strategy, we make one of them the dominant at a time, and then in a sequence we alternate between lowering the amplitude of the currently dominant transducer and increasing the amplitude of the next one, while ensuring levitation (top transducer voltage is tuned never zero). For example a sequence of; 1. Top:H, 2. S1:H and Top:L, 3. S3:H and Top:L, 4. S2:H and Top:L, 5. Top:H; leads to a 180° clock-wise rotation, where 'H' means high voltage and 'L' means low voltage. When one side-transducer is high, the next one is tuned up before the previous one is lowered down to zero. By repeating the sequence, a 360° rotation is achieved. With this, the sample is reoriented around its major axis at a step-size proportional to the change in relative voltage between the transducers and we can acquire images from each desired angle.

##### *1. Acoustic actuation by two transducers*

We found that in this (large) chamber and for the frequency used, for the elongated zebrafish embryos with a very asymmetric weight distribution (relatively heavy head and yolk) we never run into the situation where the spinning torque is larger than the restoring torque. This opens up for the alternative simpler manipulation strategy of using only the two orthogonal top and S3 transducers driven at exactly the same frequency to reorient the sample 360° around its long axis by tuning the relative phase and amplitude between them, as demonstrated in Supplementary Movie 2. In the first recording, the voltage of the top and S3 transducer is the same and is kept constant while reducing the relative phase in a step-wise manner. Rotation due to phase-modulation in a square chamber been demonstrated previously [11], where phase modulation of two degenerated ultrasonic standing modes leads to a local rotation of the pressure field and a continuous rotation of the fibre about its centre, and the rotation can be stopped at arbitrary angles (note that the fibre is not levitated in this study, but is manipulated close to the bottom of the horizontal chamber surface). Similarly in our approach, the acoustic pressure-landscape is influenced by the relative phase-shift between the two orthogonal modes which leads to a torque acting on the sample to reorient it. Each new orientation is a stable trapping position, but between each orientation (while rotating) the sample has small translations. After adjusting the phase down to $-70°$ in the second recording of Supplemenary Movie 2, we change the relative voltage between the transducers while keeping the relative phase constant, as seen in the lower recording. Starting from equal amplitudes of the two transducers, we now reduce the strength of the S3 transducer in a step-wise manner and the sample reorients to a more and more dominating top transducer.

Adjusting relative amplitude or phase, both translate to changing the pressure-landscape in the chamber, but we find that we need to adjust both to have enough degrees of freedom for rotations 360° about the samples major axis. This is in contrast to the results in Schwarz et al. [11], likely due to our step-wise modulation rather than continuous and our strongly asymmetric samples. If we adjust only the relative amplitude (at 0° relative phase) or only the phase-shift between the two transducers (at equal voltages), we can typically only sample angles between 0° to 90°, or maximum 0° to 180°. This is because of the asymmetric shape of the sample, in particular the heavy yolk sack, which determines which positions our step-wise modulation can align it to under the influence of gravity: With a dominant top transducer, the zebrafish lies flat with its (acoustically) minor axis aligned to the steepest trap-stiffness in $z$-direction, as illustrated in Supplementary Fig.3**a** with the yolk to the left or right (also seen in for 90° and 270° in Fig.2c in the main article with only one eye visible). Now, making the S3 transducer dominant will always align the zebrafish with the heavy yolk down, as illustrated in Supplementary Fig.3**a** (also seen in for 0° in Fig.2c in the main article), and alternating between dominant top and S3 only leads to re-orientations between these two end-points. However, by adjusting the relative phase in addition, we can choose which direction to rotate the sample and thus also reorient the zebrafish to orientations with the yolk facing upwards by the combination of amplitude and phase change. The working principle of acoustic reorientation combined with OCT scanning is illustrated in an animation in Supplementary Movie 1 (not to scale), created with Blender [12].

For the more spherical melanoma spheroids, shown in Supplementary Fig.4, this manipulation strategy still works, and the melanoma spheroid is reoriented around its major axis to new stable trapping orientations (sub-image *b* and Supplementary Movie 3). In Supplementary Fig.4*b* 12 angles are shown, but a more precise angle step-size is possible. For this sample certain relative phase-shift and voltage values between the two transducers lead to a sustained rotation as shown in Supplementary Fig.4**c** and second recording in Supplementary Movie 3., with rotation rate of 0.08 Hz (12.5 s period). At settings inducing a spinning torque, we can change the relative phase-shift by 180° and change the rotation direction (third recording in Supplementary Movie 3), as expected for the underlying mechanism by the spinning torque generated by exciting two orthogonal modes [10]. Note, the bright spots are accumulated dust from previous imaging which work well as markers to see the rotation direction as the image quality of the long working distance 5× objective with darkfield imaging performed here is not optimal (and much worse than for OCT imaging, as seen in Fig.3 in the main article). After changing the relative phase, the spheroid undergoes a small translational motion due to the changed force landscape, and we tune the relative voltage slightly to again have a dominating spinning torque, now with rotation in the opposite direction. This example also demonstrates that the spinning torque is easily dominated by the usually larger restoring torque for typical (slightly asymmetric) biological samples, due to the torques' different scaling laws [13]. For the current application of inducing transient rotations, these settings need to be avoided by always operating one transducer at a higher voltage and hence generating a dominating restoring torque. This latter two-transducer approach is more efficient than the four transducer approach, and opens up the possibility for simpler devices in the future, where one potentially only would need two orthogonal transducers to transiently rotate a sample. In such a chamber, the cross-section could be square and the height of the chamber would be down-scaled by a factor of two, potentially increasing the quality factor of the acoustic resonator, while still keeping the sample far away from reflecting surfaces (bottom coverslip and top transducer) that can interfere with OCT imaging. While acoustic streaming was not observed to be a problem in the current chambers, care needs to be taken when tuning the chamber dimensions and changing the sample, since the chamber dimensions and geometry influence the magnitude of acoustic streaming [14].

#### F. Reconstruction

##### 1. Data-rescaling

After data-preprocessing we obtain OCM volumes for each view. As the experiment was performed in water, we shrunk the obtained B-scans by an amount equal to the refractive index of water ($n_{\text{water}} = 1.33$) along the optical axis.

##### 2. Surface detection

After shrinking, for each volume we calculated masks containing the locations under the first surface of the specimen. This was achieved on the 3x down-sampled data by first applying a Gaussian filter (kernel size 2 pixels) followed by thresholding at the mean value of the OCM volume and summation along the optical axis with an additional thresholding step. The resulting masks were inverted and saved as Boolean, and therefore have only minor implications for memory consumption.

##### 3. Gradient-based algorithm

In the reconstruction process we found it essential to introduce the reflectivity as an optimization parameter, since the success of the alignment between volumes strongly relies on the regularization imposed on $R$. Additionally, we introduce attenuation $\alpha$ and refractive index $\Delta n$ to be able to fuse the different OCM volumes in a consistent manner, while also extracting physical and interpretable information.

The reconstruction of the maps for reflectivity $R$, attenuation $\alpha$ and refractive index $n$ are discretized on a 3-dimensional grid with Cartesian coordinates $(x, y, z)$. The OCM image formation model follows a layer-by-layer based propagation approach, where the rays are step-wise propagated through the maps for $\alpha$ and $n$ according to

$$I_{v,i}(R, \alpha, \Delta n, q_v, t_v) = T(x_i) \cdot H(x_i) \cdot R(\boldsymbol{r}_i(q_v, t_v)) \cdot \exp(-2 \sum_{j=0}^{i} \alpha(\boldsymbol{r}_j(q_v, t_v)) \Delta x_j) \quad (1)$$

$$(x_{i+1}, y_{i+1}, z_{i+1}) = (x_i + \frac{\Delta x}{1 + \Delta n(\boldsymbol{r}_i(q_v, t_v))}, y_i, z_i) \quad (2)$$

with the index $v$ describing the view, and $i$ describing the $i$th layer of the observed OCM volume. The propagation is assumed to not deviate from straight lines in the $x$ direction, since the RI of specimen is almost matched with its surrounding medium. The confocal point spread function (PSF) $T$ and roll-off $H$ are given by

$$T(x) = \frac{1}{1 + \left(\frac{x - x_{\mathrm{F}}}{2n_0 x_{\mathrm{R}}}\right)^2} \tag{3}$$

$$H(x) = \mathrm{sinc}^2\left(\frac{x - x_{\mathrm{C}}}{2x_{\mathrm{D}}}\right) \cdot \exp\left[-\frac{r}{2\ln(2)}\left(\pi \frac{x - x_{\mathrm{C}}}{2x_{\mathrm{D}}}\right)^2\right] \tag{4}$$

with $x_{\mathrm{f}}$ denoting the focal position of the beam, $2n_0 x_{\mathrm{R}}$ the depth of focus, $x_{\mathrm{C}}$ the zero-delay, $x_D$ the maximum imaging depth achievable and $r$ the ratio of optical resolution to spectral sampling pitch [15]. In our case the confocal PSF $T$ dominates compared to $H$.

The objective function of the optimization problem can be written in the following way

$$\begin{aligned} \underset{R,\alpha,\Delta n,\boldsymbol{q},\boldsymbol{t},\boldsymbol{x}_{\mathrm{F}}}{\mathrm{argmin}} \sum_v \Big[ & ||I(R, \alpha, \Delta n, q_v, t_v, x_{\mathrm{F},v}) - I_v^{\mathrm{M}}||_2^2 + \\ & \lambda_{R,\mathrm{l}_2}||M_v R||_2^2 + \lambda_{\alpha,\mathrm{l}_2}||M_v \alpha||_2^2 + \lambda_{\Delta n,\mathrm{l}_2}||M_v \Delta n||_2^2 \Big] \\ & + \lambda_{R,\mathrm{TV}}||R||_{\mathrm{TV}} + \lambda_{\alpha,\mathrm{TV}}||\alpha||_{\mathrm{TV}} + \lambda_{\Delta n,\mathrm{Tik}}||D\Delta n||_2^2, \end{aligned} \tag{5}$$

where $I_v^{\mathrm{M}}$ denotes the $v$th OCM volume, $\lambda_{p,\mathrm{TV}}$ and $\lambda_{p,\mathrm{l2}}$ and $\lambda_{\Delta n,\mathrm{Tik}}$ denote the regularization strengths on the optimization parameters. The quantities $M_v$ represent the masks which encode the first surface along the propagation direction in the OCM volumes. Furthermore, we enforce positivity on $R$, $\alpha$ and $\Delta n$. We minimize Eq. 5 using stochastic gradient descent with an diminishing step-size. For the optimization of the rotation parameters we use a reduced representation of the unit quaternions [16]. Furthermore, we also include the parameter $x_{\mathrm{F}}$ into the pool of optimization parameters, since it is not known precisely.

We compute the gradients of Eq. 5 equivalently to the forward model by propagating backwards through the volume using the vector-Jacobian product. We store the the exponential factor of Eq. 1 and the $x_i$ from Eq. 2 during the forward pass. We found this simple method to reduce the memory requirements and execution speed by a factor of $5 - 7$ when executing on the GPU.

#### III. SUPPLEMENTARY MOVIE LIST

**Supplementary Movie 1.:** Animation showing working principle of ULTIMA-OCT (not to scale), rendered in Blender[12].

**Supplementary Movie 2.:** Recording of reorientation of 3 dpf zebrafish (darkfield imaging) with two transducers by changing voltage and pressure in a step-wise manner.

**Supplementary Movie 3.:** Recording of melanoma spheroid manipulated by two transducers: stably trapped at 4 orientations, under sustained rotation, and change of rotation direction (darkfield imaging).

**Supplementary Movie 4.:** 3D rendering of a 3 dpf $Mitfa^{b692/b692}$ / $ednrb1^{b140/b140}$ zebrafish embryo showing shadow artifacts. Scale bar: 300 $\mu$m.

**Supplementary Movie 5.:** Visualization of three dimensional renderings of raw OCM volumes and reconstruction.

**Supplementary Movie 6.:** Flythroughs through raw OCM volumes and reconstruction across three axes.
